# Novel Strategy against AML by Rescuing Loss of Phase Separation of Pathogenic NPM1c

**DOI:** 10.1101/2025.04.08.646961

**Authors:** Yuting Zhu, Weifan Xu, Rong Wang, Lijuan Hu, Xiaojun Huang, Honghu Zhu, Pilong Li

## Abstract

Nucleophosmin (NPM1) is a nucleolar protein that participates in many biological processes through phase separation. In acute myeloid leukemia (AML), nearly one-third of cases harbor *NPM1c* mutations. Previous studies mainly focused on the contribution of the additional NES in NPM1c mislocalization and AML development; however, the mechanisms underlying the relationship between aberrant localization and phase separation of NPM1c remain poorly understood. In this study, we demonstrate that loss of phase separation (LoPS) with RNA is associated with NPM1c mislocalization, which uncovers a new strategy for therapeutic targeting of NPM1c. Leveraging high-throughput screening, we identify a potent small molecule capable of rescuing NPM1c LoPS *in vitro* and effectively restoring the nucleolar localization of NPM1c within cells. Remarkably, this compound demonstrates anti-leukemia activity in both AML cell lines and patient-derived blasts. Overall, these findings indicate a promising avenue for developing new AML therapies against NPM1c by reinstating its phase separation capabilities.

**One-Sentence Summary:** Rescuing the lost phase separation capacity of NPM1c unveils a promising AML therapy.

## Introduction

Acute myeloid leukemia (AML) is a kind of hematologic malignancy characterized by abnormal differentiation and excessive proliferation of hematopoietic stem cells. In recent decades, the primary treatment approach for AML is the “3+7 regimen” involving the combination of anthracyclines and cytarabine. However, due to the strong heterogeneity of the disease, relapse remains a considerable challenge^1,2^. Currently, novel targeted therapies are under investigation as potential alternatives in order to achieve higher cure rates^3^.

Within the landscape of AML, one of the most prevalent and intriguing mutations is located within the *NPM1* gene. It accounts for approximately one-third of newly diagnosed cases^4^. NPM1 consists of an N-terminal oligomeric domain, a central intrinsically disordered region (IDR), and a C-terminal RNA-binding domain (RBD)^5^. NPM1 is a multifunctional nucleolar protein involved in various physiological processes, such as ribosome synthesis, chromatin remodeling and DNA repair^6^. The *NPM1* mutation (*NPM1c*), characterized by 4-bp insertions in exon 12, produces a protein with a newly generated nuclear export signal (NES) and cytoplasmic mislocalization^7^. NPM1c plays a crucial role in both the initiation and progression of AML^8,9^, suggesting its potential as a possible therapeutic target. Numerous studies have focused on investigating the pathogenic mechanisms of NPM1c. On one hand, the cytoplasmic translocation of NPM1c results in nuclear export of numerous tumor suppressors and transcription factors, which contributes to AML progression indirectly^10–12^. On the other hand, nuclear NPM1c is a direct gene activator regulating the expression of oncogenes through its chromatin binding^13–15^. These studies consistently demonstrate the significance of NPM1c mislocalization.

Several strategies have been investigated for targeting NPM1c in AML treatment^16–18^. Combined treatment with all-trans retinoic acid (ATRA) and arsenic trioxide (ATO) has shown the ability to specifically degrade the NPM1c protein, leading to apoptosis and promoting cellular differentiation^19^. The small molecule NSC348884 selectively targets the oligomerization domain of NPM1, inhibiting NPM1 multimerization and subsequently triggering apoptosis^20^. Chimeric antigen receptor (CAR) T-cell therapy, tailored to target NPM1c epitopes, effectively eradicates NPM1c^+^cells^21^. Other approaches involve reducing *HOX*/*MEIS1* gene expression, either by correcting the localization of NPM1c with the XPO1 inhibitor Selinexor^22^, or using MLL1–Menin inhibitors^15,23,24^.

Despite these encouraging avenues, there is currently a lack of clinically approved drugs specifically targeting NPM1 mutants to alleviate AML. Intriguingly, in addition to generating a new NES, *NPM1c* mutations significantly impair the structure of the RBD, which is essential for the heterotypic phase separation of NPM1. Phase separation is crucial for NPM1’s nucleolar localization and biological functions^25–29^. This indicates a potential relationship between aberrant phase separation of NPM1c and its pathogenic characteristics.

Here, we present a new perspective on NPM1c by examining it through the lens of phase separation. Our findings uncover a significant deficiency in NPM1c phase separation capability, resulting in the formation of cytoplasmic aggregates. We further demonstrate that loss of phase separation (LoPS) with RNA is closely associated with the abnormal cellular localization of NPM1c. Through *in vitro* high-throughput screening, we identify compounds capable of rescuing NPM1c’s LoPS and we succeed in finding a candidate compound that restores NPM1c’s nucleolar localization. Significantly, this compound also exhibits anti-leukemia activity in AML cell lines and patient-derived blasts. Overall, these findings underscore the therapeutic potential of targeting the aberrant phase separation of NPM1c.

## Results

### NPM1 Exhibits RNA-Dependent Phase Separation and Nucleolar Localization

Previous studies have characterized the liquid-liquid phase separation (LLPS) of NPM1 and rRNA, which occurs through heterotypic interactions^25,30–32^. We found that overexpressed mCherry-NPM1 predominantly localized within the nucleolus and exhibited liquid-like behavior as expected (**Figures1A and 1B**). Consistent with previous studies, recombinant NPM1 protein formed spherical liquid-like droplets in the presence of RNA *in vitro*, and demonstrated fluidity similar to that observed in cells. (**Figures1C and 1D**). Notably, the size of *in vitro*-formed droplets is positively correlated with NPM1 and RNA concentrations, suggesting that RNA promotes NPM1 phase separation. We further investigated the role of RNA in NPM1 phase separation and nucleolar localization. RNaseA, an endoribonuclease with specific RNA-digesting capabilities, induced the translocation of approximately 70% of NPM1 from the nucleolus to the cytoplasm in Hela cells (**Figures1E and S1A**). Accordingly, *in vitro*-formed recombinant NPM1 droplets disappeared within 2 minutes of adding RNaseA (**Figure1F**). Collectively, our findings from both *in cellulo* and *in vitro* experiments unequivocally confirm the intricate relationship between NPM1’s nuclear localization and its heterotypic LLPS with RNA.

**Figure1.**
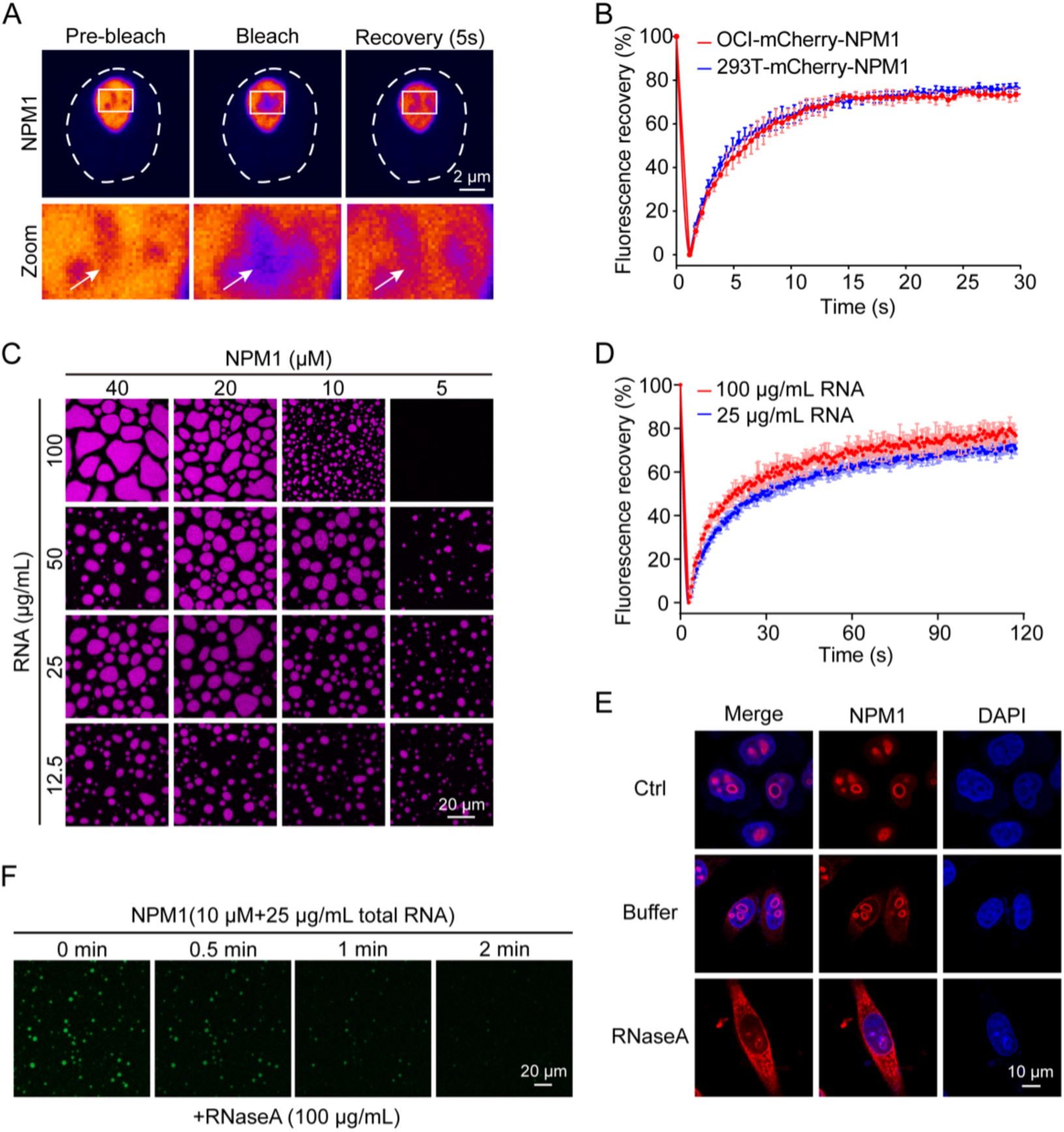
Phase separation and nucleolar localization of NPM1 rely on RNA. **(A)** Representative image of the FRAP of mCherry-NPM1 in OCI-AML3 cells, with the nucleus marked by white dashed lines. **(B)** FRAP quantification for overexpressed mCherry-NPM1 in HEK293T and OCI-AML3 cells, with n=5 replicates. **(C)** Phase diagram illustrates the phase separation between NPM1 and RNA *in vitro*. NPM1 is labeled with Alexa561 dye. **(D)** FRAP analysis demonstrates the dynamic behavior of condensates formed by NPM1 (20 μM) and RNA *in vitro*, with n=5 replicates. **(E)** Representative images of immunofluorescence showing the relocation of NPM1 from the nucleolus to the cytoplasm in Hela cells following RNase A treatment. **(F)** Representative images of *in vitro*-reconstituted NPM1 condensates with RNase A treatment.

### RNA-Dependent Phase Separation and Thermostability of NPM1c is Impaired

Alterations in NPM1c result in the destabilization of the C-terminal globular domain^33,34^, while concomitantly increasing its propensity for amyloid aggregation^35,36^. We next explored the characteristics of NPM1 and NPM1c in OCI-AML3 cells, which harbor a heterozygous *NPM1c* mutation. Our nuclear and cytoplasmic extraction experiments confirmed that substantial proportions of NPM1 and NPM1c translocate into the cytoplasm (**FigureS2A**). Consistent with previous studies, immunofluorescence results revealed cytoplasmic localization of both NPM1 and NPM1c (**Figure2A**)^11,37,38^. We observed that NPM1c formed distinct bright granules in the cytoplasm, which was confirmed by immunoelectron microscopy (iEM) (**Figure2B**). The environments of the nucleolus and cytoplasm differ significantly, such as in their RNA concentration, which has been shown to regulate phase separation behavior^39^. To further investigate the dynamic state of NPM1c in the cell, we established OCI-AML3 cells stably overexpressing mEGFP-NPM1c. NPM1c is localized in both the nucleolus and the cytoplasm (**Figure2C**). In contrast to the fluid-like state of the nucleolar protein, NPM1c in the cytoplasmic granules exhibited a more solid-like character (**Figures2C and 2D**).

**Figure2.**
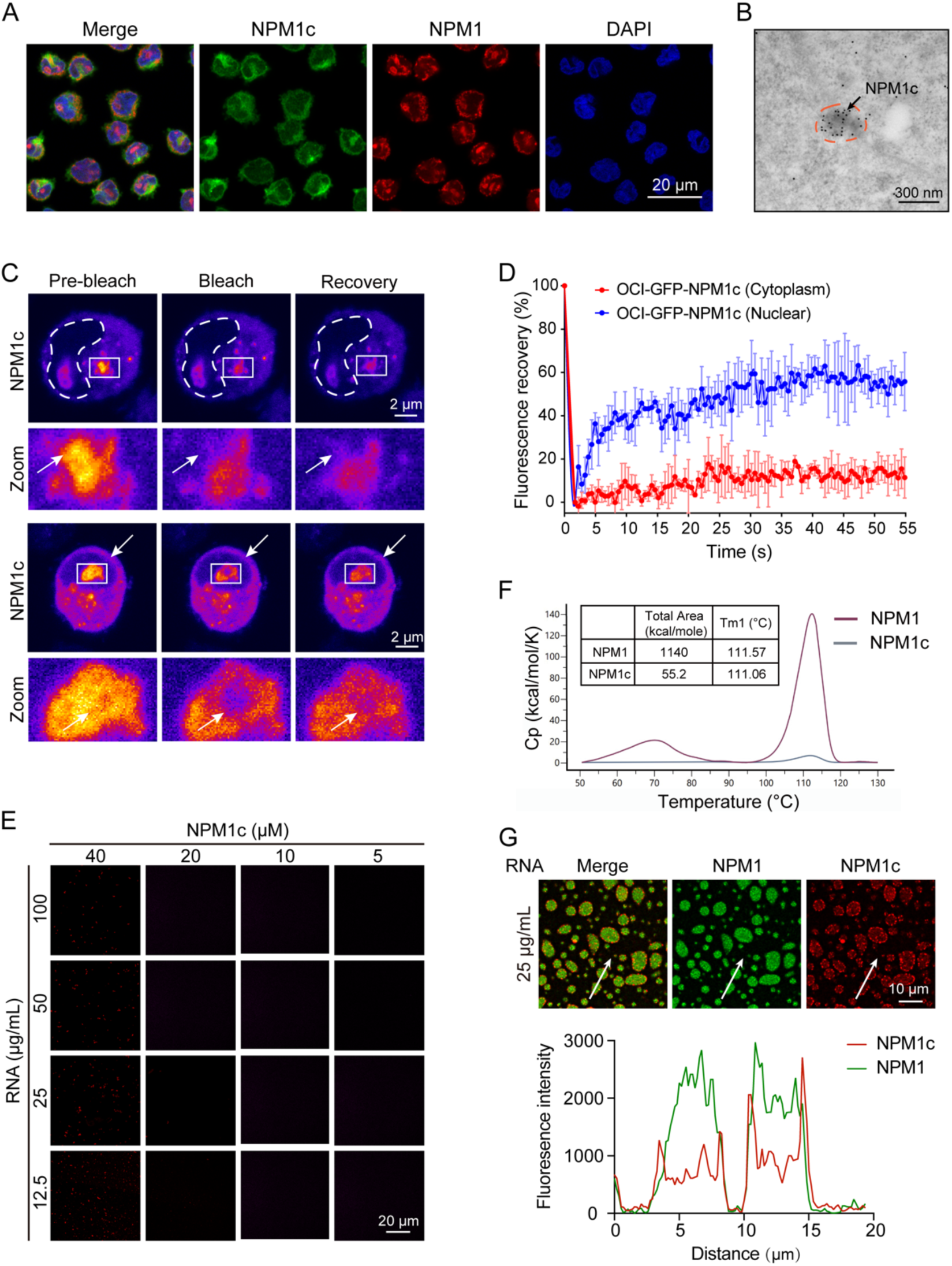
Impaired phase separation of NPM1c with RNA. **(A)** Representative immunofluorescence microscopy image shows the subcellular localization of NPM1 and NPM1c in OCI-AML3 cells. **(B)** Representative immunoelectron microscopy image shows the cytoplasmic aggregation of NPM1c. **(C)** Representative images of FRAP for overexpressed GFP-NPM1c in the cytoplasm (top; the nucleus is marked by dashed white lines) and the nucleolus (bottom) of OCI-AML3 cells. **(D)** Quantification of FRAP data in (C) demonstrates that the cytoplasm-localized NPM1c exhibits an elevated tendency for aggregation. n=5. **(E)** Phase diagram illustrates that NPM1c loses the capacity for phase separation with RNA *in vitro*. NPM1c is labeled with Alexa647 dye. **(F)** Differential scanning calorimetry (DSC) results of purified NPM1 and NPM1c proteins. T_m_ represents the transition midpoint. Total area represents enthalpy (ΔH), which indicates the total heat energy uptake by the sample. Cp represents heat capacity. **(G)** Top: representative image of the reconstituted NPM1/NPM1c/RNA three-component phase separation system. Bottom: colocalization analysis between NPM1 and NPM1c. Fluorescence intensity of NPM1 and NPM1c is plotted along the white arrow. NPM1 is labeled with Alexa561 dye; NPM1c is labeled with Alexa647 dye.

The concentrations of endogenous NPM1 and NPM1c in OCI-AML3 cells were determined to be 12.8 μM and 10.9 μM respectively, exceeding the saturation concentration (Csat) for NPM1 phase separation (**FigureS2B**). The NPM1 concentration in HL-60 cells was 23.4 μM, which was approximately equal to the combined concentration of NPM1 and NPM1c in OCI-AML3 cells. Thus, the presence of a heterozygous mutation reduces the concentration of wild-type protein. We performed the *in vitro* phase separation assay for NPM1c under physiological conditions, as used for NPM1. In contrast to the robust phase separation capability of NPM1 (**Figure1C**), almost no condensate formation was observed for NPM1c, indicating a deficiency in its ability to undergo heterotypic phase separation with RNA (**Figure2E**). Given that NPM1 is also known to undergo homotypic phase separation^30,40^, we investigated the LLPS potential of NPM1 and NPM1c in the presence of a crowding agent. Although both proteins formed spherical droplets, NPM1c exhibited a lower propensity for phase separation (**FigureS2C**). Furthermore, FRAP analysis revealed that NPM1 condensates were dynamic with a high recovery rate, whereas the recovery of NPM1c condensates was negligible (**FigureS2D**), which is consistent with the solid-like property of NPM1c condensates observed in the cytoplasm.

Differential scanning calorimetry (DSC) results further indicated that NPM1 requires a higher enthalpy (ΔH) than NPM1c to disrupt the tertiary structure (**Figure2F**). The DSC data suggest that NPM1 possesses a more intact structure and greater thermostability. When exposed to hyperthermia (50°C, 30 minutes), *in vitro*-formed NPM1c homotypic phase-separated condensates were destabilized, appearing as bead-like structures, while the morphology of NPM1 homotypic condensates were unaffected (**FigureS2E**). This observation suggests that NPM1c condensates are more sensitive to thermal changes than NPM1 condensates. Thermostable proteins exhibit reduced susceptibility to thermal aggregation than unstable proteins^41^. When OCI-AML3 cells were exposed to mild hyperthermia (41°C), NPM1 remained largely soluble while NPM1c was robustly destabilized, leading to its insolubility (**FigureS2F**).

Collectively, these data suggest that NPM1c lacks the ability to undergo phase separation with RNA. Although NPM1c retains some of its homotypic phase separation capability, its characteristics including Csat and dynamic behavior are different from those of NPM1. Moreover, NPM1c tends to aggregate and is more susceptible to hyperthermia-induced destabilization.

### LoPS is the Underlying Mechanism for the Mislocalization of NPM1c

NPM1c-positive AML patients always harbor both NPM1 and NPM1c, suggesting the presence of a three-component phase separation system involving NPM1, NPM1c and RNA. To further investigate the characteristics of this system, we co-transfected mCherry-NPM1 and mEGFP-NPM1c into HEK293T cells and subsequently confirmed their interaction through co-immunoprecipitation (**FigureS3A**). In contrast to OCI-AML3 cells (**Figures2C and 2D**), both proteins were evenly distributed in the cytoplasm, with no visible condensate formation (**FigureS3B**). This suggests that overexpressed NPM1 might possess the ability to disperse NPM1c aggregates in the cytoplasm through N-terminus-mediated heterotypic interactions.

Next, we mixed NPM1 and NPM1c at a concentration of 10 μM in the presence of RNA to form condensates *in vitro*. NPM1 was enriched within the droplets while NPM1c adhered to the droplet surface. Thus, NPM1 and NPM1c showed a mutually exclusive distribution in the condensates (**Figure2G**). The centrifugation assay further confirmed that the partitioning of NPM1 into droplets was positively correlated with RNA concentrations, while NPM1c was consistently excluded from condensates (**FigureS3C**). Molecular interaction landscapes can be drastically altered in phase-separated condensates versus in dilute solutions^42^. Consistently, our results indicate that the heterotypic interactions between NPM1 and NPM1c, presumably via the N-terminal oligomerization domain, are compromised by the forces generated by NPM1 and RNA phase separation, leading to the exclusion of NPM1c from NPM1/RNA condensates. Given that NPM1 functions as a scaffold protein within the nucleolus^25,28^, these findings prompt us to consider whether NPM1c’s mislocalization in AML patient cells is associated with its LoPS with RNA. Within the reconstituted three-component condensates, NPM1c exhibited enhanced mobility compared to homotypic phase separation conditions (**FiguresS3D and S2D**), aligning with observations that nucleolar NPM1c possesses higher mobility compared to cytoplasmic NPM1c (**Figures2C and 2D**).

We further investigated the role of phase separation in NPM1c nucleolar localization. The RBD is indispensable for NPM1 phase separation^25^. Amino acids Trp^288^ and Trp^290^, which are located in the hydrophobic core of the NPM1 RBD, play a crucial role in maintaining the integrity of the RBD structure and the nucleolar localization of NPM1^34,43^. Notably, Trp^288^ and Trp^290^ are mutated in NPM1c. Therefore, we reintroduced Trp^288^ and Trp^290^ into NPM1c and generated a mutant named NPM1c^288/290W^ (**Figures3A**). Intriguingly, NPM1c^288/290W^ was found to have the capability of forming droplets and exhibited perfect relocation to the nucleolus (**Figures3B and 3C**). The centrifugation assay also confirmed the phase separation capacity of NPM1c^288/290W^ with RNA (**Figure3D**). In the NPM1/NPM1c^288/290W^/RNA three-component phase separation system, NPM1c^288/290W^ readily distributed into NPM1 droplets (**Figure3E**). We replaced the RBD of NPM1c with a RBDs from a variety of different proteins: nucleolar proteins nucleolin (NCL) and fibrillarin (FBL), and cytoplasmic proteins YTHDF1 and CIRBP. Three out of four chimeric proteins showed nucleolar localization, while the RBD-only constructs only rarely exhibited nucleolar enrichment (**Figure3F**).

**Figure3.**
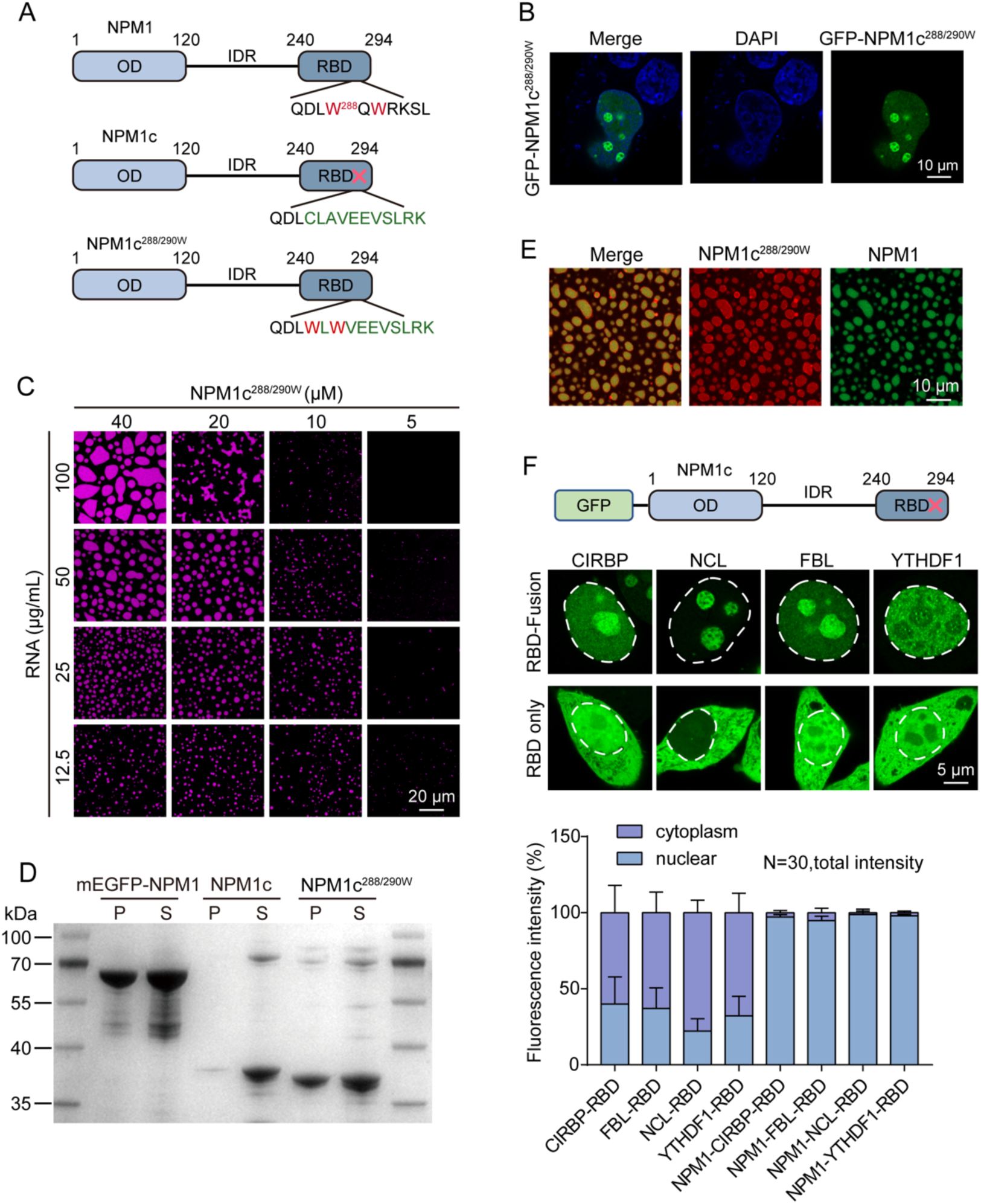
Restoration of NPM1c’s phase separation ability and nucleolar localization by repairing its RBD. **(A)** Schematic representation of the NPM1, NPM1c and NPM1c^288/290W^ constructs used in this study. The N-terminal oligomeric domain (OD), an intrinsically disordered region (IDR), and a C-terminal RNA-binding domain (RBD) are depicted. Amino acids changes of NPM1c are indicated in green, the 288W and 290 W of NPM1c^288/290W^ are indicated in red. **(B)** Representative immunofluorescence microscopy image shows the localization of overexpressed GFP-NPM1c^288/290W^ in HEK293T cells. **(C)** Phase diagram of NPM1c^288/290W^ and RNA *in vitro*. NPM1c^288/290W^ is labeled with Alexa647 dye. **(D)** Centrifugation assay demonstrates the enrichment of NPM1, NPM1c and NPM1c^288/290W^ in the *in vitro* phase separation system, where P indicates the pellet and S represents the supernatant. Protein concentration is 10 μM, RNA concentration is 100 μg/mL. **(E)** Representative image of the *in vitro*-reconstituted NPM1/NPM1c/ RNA three-component phase separation system. NPM1 is labeled with Alexa561 dye; NPM1c is labeled with Alexa647 dye. Protein concentration is 10 μM, RNA concentration is 100 μg/mL. **(F)** Top: a schematic diagram of the GFP-tagged NPM1c protein, with its defective RBD. Chimeric proteins were created by replacing the NPM1c RBD with the RBD of four other proteins. Middle: images showing the localization of overexpressed GFP-RBD and GFP-NPM1c-RBD fusion proteins in HEK293T cells. Bottom: statistical results of the average fluorescence intensity of the indicated fusion proteins (GFP-RBD only and GFP-NPM1c-RBD) in the cytoplasm and nucleus, with n=30.

Overall, these results underscore the significance of LLPS in NPM1c localization, and suggest that restoring phase separation with RNA could potentially restore NPM1c’s nucleolar localization. This might be a promising strategy for AML treatment.

### High-Throughput Screening Identifies Compounds that Rescue NPM1c LoPS

We hypothesized that the ability of NPM1c to undergo LLPS *in vitro* may partially reflect its cellular phase separation ability and nucleolar localization. Hence we attempted to identify compounds capable of restoring the ability of NPM1c to undergo phase separation in the presence of RNA. To this end, we conducted a high-throughput screen of 2,572 compounds from a library of FDA-approved drugs. The *in vitro* screen was conducted in the presence of 150 mM NaCl, 20 μM Alexa488 labeled NPM1c, and 25 μg/mL RNA. Compounds at a final concentration of 50 μM were added to the pre-mix of NPM1c and RNA, followed by high-content microscopy imaging (**Figure4A**). We used the partition coefficient of NPM1c as the evaluation index: a higher fluorescence intensity within droplets (“in signal”) compared to outside droplets (“out signal”) indicates a greater enrichment of NPM1c in droplets and a higher degree of phase separation. Among the top-ranking compounds, only Luteolin, Apigenin, and Lenvatinib induced the formation of spherical NPM1c condensates in the presence of RNA (**Figures4B and 4C**). Other compounds induced irregularly shaped NPM1c-positive structures (**FigureS4A**). ATP has been proved to interact with NPM1 and promote phase separation between RNA and NPM1, and increase the partitioning of both components^31^. Notably, Apigenin and Lenvatinib are ATP-competitive inhibitors^44,45^, and Luteolin possesses a similar structure to Apigenin (**FigureS4B**), which suggests that these compounds may function in a similar way.

**Figure4.**
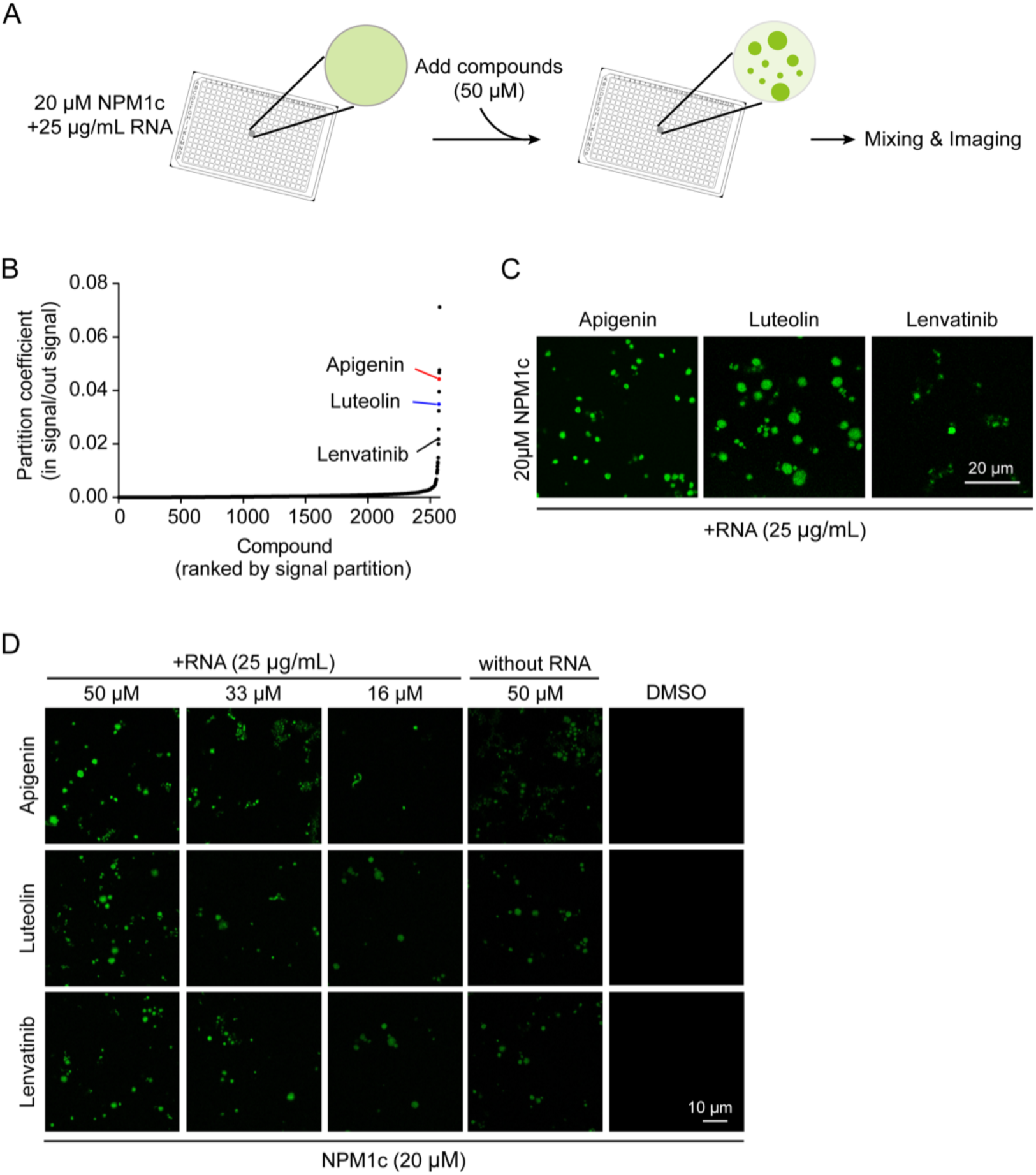
High-throughput screening for compounds that restore the phase separation of NPM1c. **(A)** Workflow of the *in vitro* screening assay for compounds rescuing LoPS of NPM1c. Alexa488 labeled NPM1c (20 μM) and RNA (25 μg/mL) were premixed, then compounds at a final concentration of 50 μM were added to the system, fllowed by high-content microscopy imaging. **(B)** Compounds are ranked by the partitioning of the GFP signal within NPM1c droplets. Apigenin, Luteolin and Lenvatinib are highlighted. **(C)** Representative images illustrate examples of compounds (50 µM) leading to the robust partitioning of NPM1c into droplets *in vitro*. RNA concentration is 25 μg/mL. **(D)** NPM1c (20 μM) concensates induced by different concentrations of the identified compounds. Compound concentration is labeled at the top of the figure (50 µM, 33 µM, 16 µM), RNA concentration is 25 μg/mL. In (C) and (D), NPM1c is labeled with Alexa488 dye.

We investigated the effects of Luteolin, Apigenin, and Lenvatinib on NPM1c phase separation in more detail. The compounds were found to induce condensation of NPM1c in the absence of RNA, while RNA further enhanced the phase separation of NPM1c (**Figure4D**). This observation indicates that RNA also participates in the phase separation process. Additionally, we observed a positive correlation between the number and size of NPM1c droplets and compound concentration. FRAP analysis suggested that NPM1c droplets induced by Lenvatinib exhibit low fluidity, which is similar to homotypic NPM1c condensates (**FigureS4C**). For NPM1, treatment with the three different compounds led to the formation of new condensates with varying morphology and fluorescence intensity (**FigureS4D**). Thus, these compounds also alter the phase separation of NPM1 to some extent.

### Lenvatinib Reverses NPM1c Localization and Exhibits Antineoplastic Activity

We further explored whether these compounds can impact the subcellular localization of endogenous NPM1 and NPM1c proteins in OCI-AML3 cells. Interestingly, Lenvatinib caused the nuclear relocation of both NPM1 and NPM1c, while Apigenin and Luteolin did not show strong effects (**Figure5A**). Additionally, Lenvatinib functions in a time- and concentration-dependent manner (**FiguresS5A and S5B**). To further validate the impact of Lenvatinib on NPM1c translocation, we treated HEK293T cells stably expressing mEGFP-NPM1c with Lenvatinib. Live-cell imaging revealed that Lenvatinib led to partial nucleolar relocation of NPM1c (**Figures5B and S5C**). Moreover, Lenvatinib also induced nucleolar re-localization of NPM1c in primary AML cells from patients and enhanced the Pearson correlation coefficient (PCC) of NPM1 and NPM1c (**Figures 5D and S5D**). These results suggest that Lenvatinib efficiently restores nucleolar localization of NPM1c.

**Figure5.**
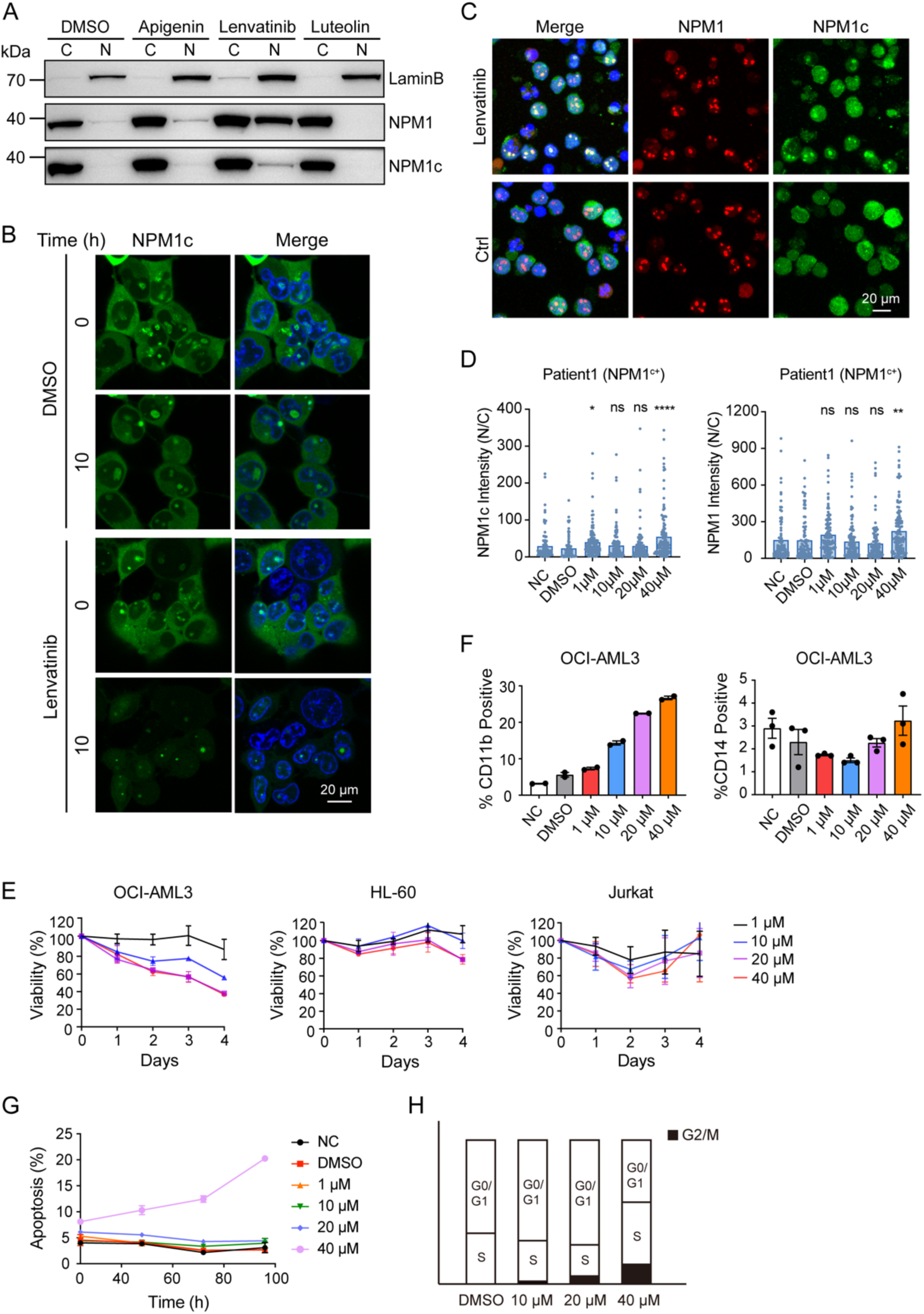
Lenvatinib reverses abnormal NPM1c localization and exhibits anti-leukemia activity. **(A)** Western blot analysis of NPM1 and NPM1c levels in the nucleus and cytoplasm of OCI-AML3 cells after 24-hour treatment with the indicated compounds. “C” indicates the cytoplasm, “N” represents the nucleus, and LaminB serves as the marker protein for the nucleus. **(B)** Representative live-cell images showing the localization of overexpressed mEGFP-NPM1c in HEK293T cells after 10-hour Lenvatinib treatment (40 µM) in HEK293T cells. **(C)** Representative immunofluorescence images of endogenous NPM1 and NPM1c localization upon 24-hour Lenvatinib treatment (40 µM) in patient-derived blasts harboring a heterozygous *NPM1c* mutation. **(D)** Statistical analysis of nuclear intensity/cytoplasmic intensity (N/C) of NPM1 and NPM1c in the presence of increasing concentrations of Lenvatinib. **(E)** Relative viability of OCI-AML3, HL-60 and Jurkat cells following 24, 48, 72, 96-hour treatment with Lenvatinib at the indicated concentrations. **(F-H)** Differentiation (F), apoptosis (G) and cell cycle analysis (H) in OCI-AML3 cells exposed to the indicated concentrations of Lenvatinib. Lenvatinib treatment lasted for 96 hours in the differentiation assay, 24, 48, 72, 96 hours in the apoptosis assay, and 24 hours for the cell cycle assay.

We further investigated the anti-leukemia activity of Lenvatinib. The anti-proliferative activity of Lenvatinib was assessed in different leukemia cell lines, with OCI-AML3 showing the highest sensitivity to Lenvatinib treatment. In contrast, Lenvatinib exerted a weak effect on HL-60 and Jurkat cells, which do not carry the *NPM1c* mutation (**Figure5E**). Additionally, we observed that Lenvatinib triggered the dose-dependent differentiation, apoptosis, and proliferation of OCI-AML3 cells (**Figures 5F and 5G-H**). In summary, these results demonstrate that Lenvatinib possesses anti-leukemia activity in NPM1c^+^ AML cell lines and patient-derived blasts, highlighting its therapeutic potential for targeting NPM1c in AML.

## Discussion

Due to its high frequency in AML, the NPM1c mutant protein is considered an ideal target for AML therapy. Several strategies targeting NPM1c, such as XPO1 inhibitors and Menin inhibitors are currently undergoing early clinical trials^46,47^. However, these strategies do not specifically target NPM1c, and therefore they may cause unwanted side effects. For example, XPO1 is a broad-spectrum nuclear receptor, which means that the XPO1 inhibitor Selinexor also mediates nuclear retention of a variety of tumor suppressor factors such as TP53 and pRB. Selinexor thus exhibits low specificity and high toxicity. Therefore, new drugs that can effectively regulate NPM1c nuclear-cytoplasmic translocation will provide a new therapeutic strategy for NPM1c-positive AML.

Phase separation-mediated biomolecular condensates are essential for various cellular processes, including cellular compartment formation, gene regulation, signal transduction, *etc*.^28,48,49^. Many diseases are associated with aberrant phase separation, which provides new strategic options for targeted therapy and drug discovery, especially for diseases caused by undruggable targets^50^. Disease-associated aberrant phase separation can be defined as loss of phase separation (LoPS) and gain of phase separation (GoPS). LoPS involves reduced phase separation ability, while GoPS refers to newly acquired or increased phase separation ability^51^. Current evidence illuminates the pathological consequences of both LoPS and GoPS. Although there have been some studies of drugs targeting GoPS^52–54^, strategies focusing on reversing LoPS is limited. Our investigation reveals that LoPS is an important mechanism underlying the impact of the NPM1c mutant in AML pathogenesis. NPM1 performs multiple functions depending on its phase separation properties, and the *NPM1c* mutation severely disrupts the function of the nucleic acid-binding domain, which is essential for the heterotypic phase separation of NPM1. Understanding how the *NPM1c* mutation disturbs the phase separation properties is essential for unraveling the complex pathogenic mechanisms of AML.

Our results reveal that the cytoplasmic mislocalization of NPM1c is closely linked to the LoPS between NPM1c and RNA. The nucleolar localization of wild-type NPM1 is highly dependent on its phase separation with RNA (**Figure6A**). The NPM1c mutant exhibits LoPS with RNA and is excluded from the three-component phase separation system NPM1/RNA/NPM1c, which may be the underlying mechanism of NPM1c mislocalization. Additionally, NPM1c still retains partial homotypic phase separation ability, and mislocalized NPM1c tends to form weakly dynamic aggregates through self-interaction in the cytoplasm (**Figure6B**). This suggests that LoPS of NPM1c in the nucleolus results in its GoPS in the cytoplasm, indicating that LoPS and GoPS can coexist under specific conditions. Furthermore, just two mutations in the RBD can efficiently restore NPM1c phase separation and nucleolar localization, which suggests that compounds may possess a similar ability. We further conducted an *in vitro* screen, and we identified that Lenvatinib rescues LoPS of NPM1c, restores its nucleolar localization, and exhibits anti-leukemia activity (**Figure6C**).

**Figure6.**
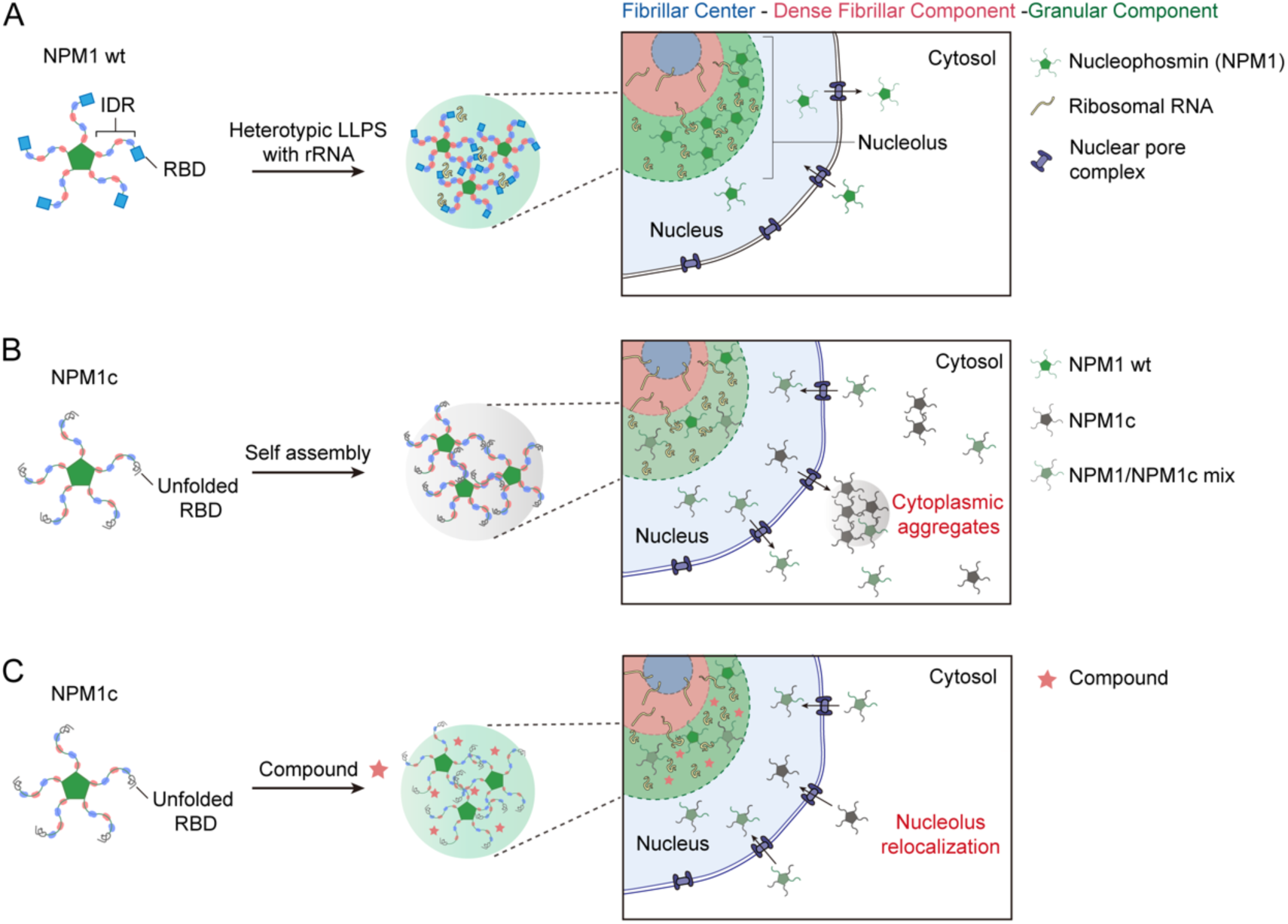
Working models showing NPM1 in normal cells, NPM1c in AML pathogenesis, and drug-induced restoration of NPM1c’s phase separation ability. **(A)** Left: Schematic diagram of NPM1 undergoing LLPS with RNA. Right: NPM1 exhibits RNA-dependent phase separation and localizes to the granular component of the nucleolus in wild-type cells. Green pentagon, OD of NPM1. **(B)** RNA-dependent phase separation is impaired for NPM1c, leading to mislocalization in the cytoplasm and formation of weakly dynamic aggregates. **(C)** Compounds can restore the correct nucleolar localization of NPM1c by rescuing its LoPS.

Overall, our findings demonstrate the therapeutic potential of targeting NPM1c’s LoPS in AML treatment, which is different from conventional GoPS-based strategies. However, it is essential to provide robust *in vivo* efficacy data and comprehensive drug chemistry structure-activity relationship (SAR) data to substantiate the potential of this new treatment paradigm.

## Data availability

All data are available from the corresponding author upon reasonable request.

## Acknowledgements

This work was supported by grants from the National Natural Science Foundation of China (32150023 to P.L., and 32100990 to W.F.). We are grateful to SLSTU-Nikon Biological Imaging Center (Center of Pharmaceutical Technology, Tsinghua University, Beijing, China) for imaging support, and Ting Wang (High Throughput Screening Core Facility, Center of Pharmaceutical Technology, Tsinghua University, Beijing, China) for High Throughput Screening assays. We also thank the Tsinghua University Branch of China National Center for Protein Sciences (Beijing) and Tsinghua University Technology Center for Protein Research for the Cell Function Analyzing Facility support.

## Author contributions

P.L.L., Y.T.Z. and W.F.X. conceived the study. Y.T.Z. and W.F.X. designed experiments. Y.T.Z. performed experiments. Y.T.Z. contributed to data analyses, interpretation and editing the manuscript. All authors reviewed and discussed the manuscript. Y.T.Z. and W.F.X. wrote the paper and P.L.L. supervised all aspects of the work and approved the final manuscript.

## Supplementary Figures & Figure Legends

**FigureS1.**
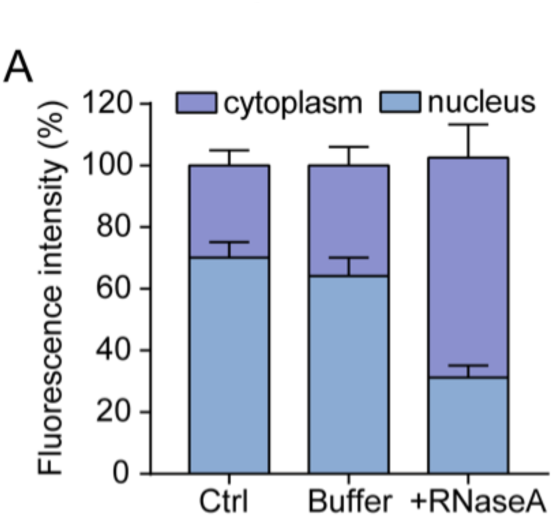
RNase A treatment disrupts the nuclear localization of NPM1. **(A)** Statistical analysis of the total fluorescence intensity of NPM1 in the nuclelus and the cytoplasm following RNase A treatment (10 µg/ml, 30 minutes), with n=30 replicates.

**FigureS2.**
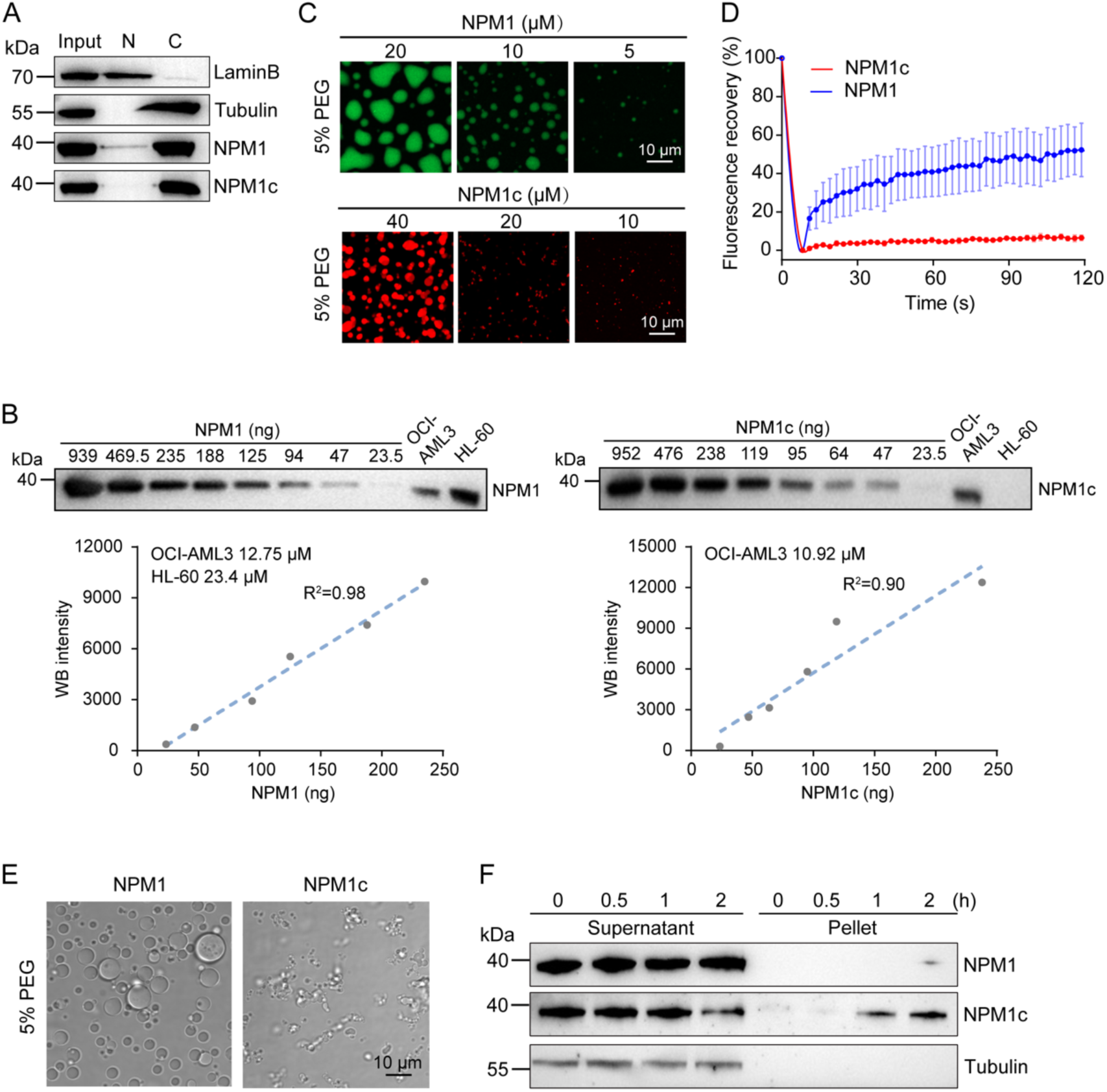
NPM1c exhibits reduced mobility and decreased thermal stability compared to wild-type NPM1. **(A)** Western blot analysis of NPM1 and NPM1c levels in the OCI-AML3 cell line, which harbors a heterozygous *NPM1c* mutation. **(B)** Quantification of NPM1 and NPM1c concentrations in OCI-AML3 cells and HL-60 cells. To generate standard curves, the indicated amounts of purified recombinant NPM1 and NPM1c proteins were detected by immunoblotting and the gray values were plotted (R^2^ = 0.98 for the NPM1 curve and 0.90 for the NPM1c curve). These curves were then used to quantify the protein levels in cell lysates (right lanes of the immunoblot). **(C)** *In vitro* homotypic phase separation of NPM1 and NPM1c in the presence of 5% PEG8000. **(D)** FRAP analysis demonstrating the dynamic behavior of *in vitro-*formed homotypic NPM1 and NPM1c droplets. **(E)** Morphological changes of *in vitro-*formed NPM1 and NPM1c condensates (40 µM of each protein) after incubation at 50°C for 30 minutes. **(F)** Western blot analysis of OCI-AML3 NPM1 and NPM1c levels in supernatant and insoluble fractions after the cells were exposed to hyperthermia treatment (42°C) for 0.5, 1, 2 hours.

**FigureS3.**
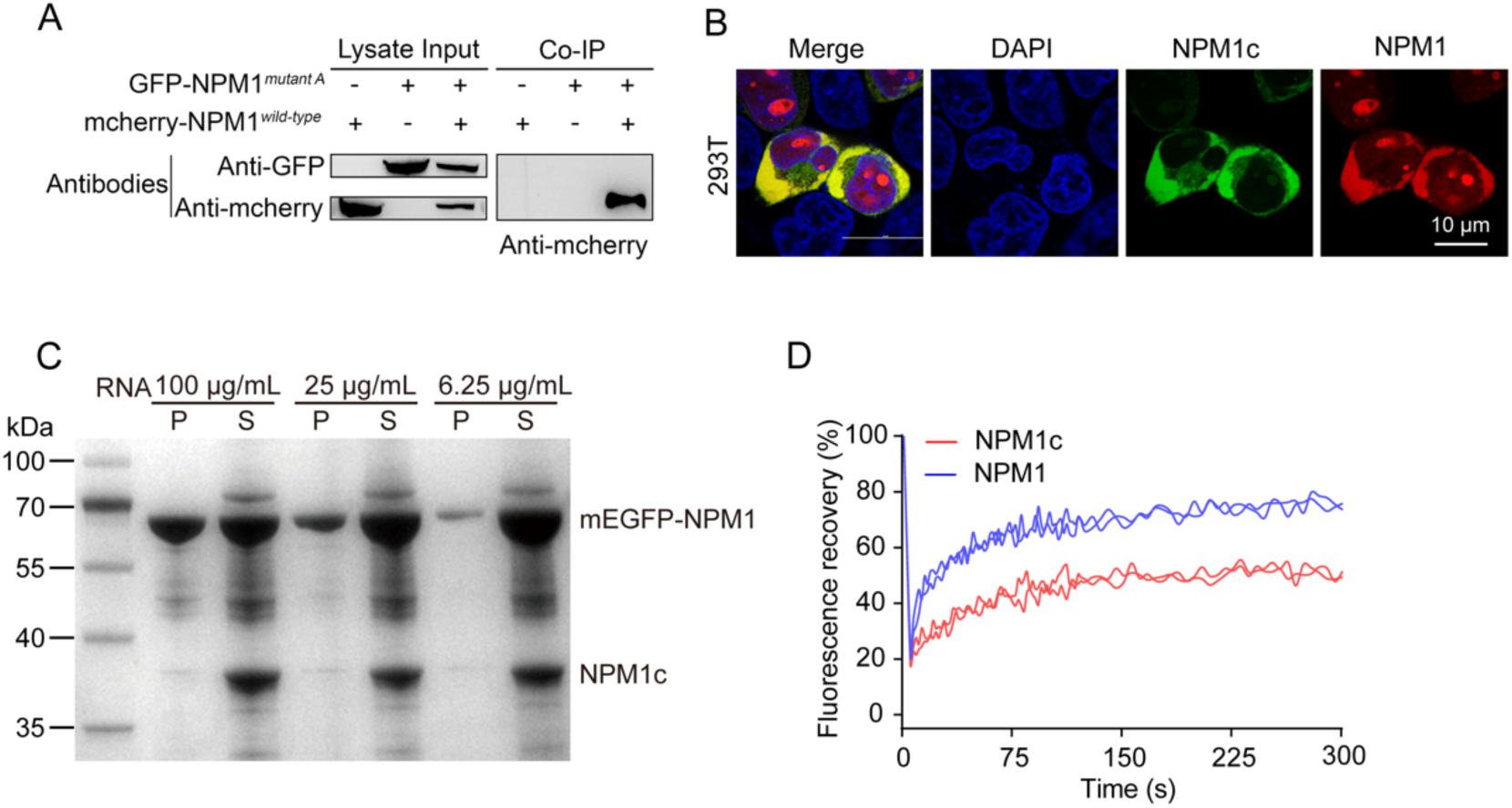
Characterization of the interaction between NPM1c and NPM1 in cells and in *in vitro*-reconstituted three-component condensates. **(A)** Co-immunoprecipitation of mEGFP-NPM1c and mCherry-NPM1 in HEK293T cells reveals an interaction between NPM1 and NPM1c. **(B)** Confocal image shows the location of overexpressed mEGFP-NPM1c and mCherry-NPM1 in HEK293T cells. **(C)** Centrifugation assay demonstrates the exclusion of NPM1c from NPM1 and RNA condensates formed *in vitro*. Protein concentrations were 10 µM. P indicates the pellet and S represents the supernatant. **(D)** FRAP analysis shows the dynamic behavior of NPM1 and NPM1c in three-component condensates formed *in vitro*. Protein concentrations were 10 µM; RNA concentration was 25 µg/mL.

**FigureS4.**
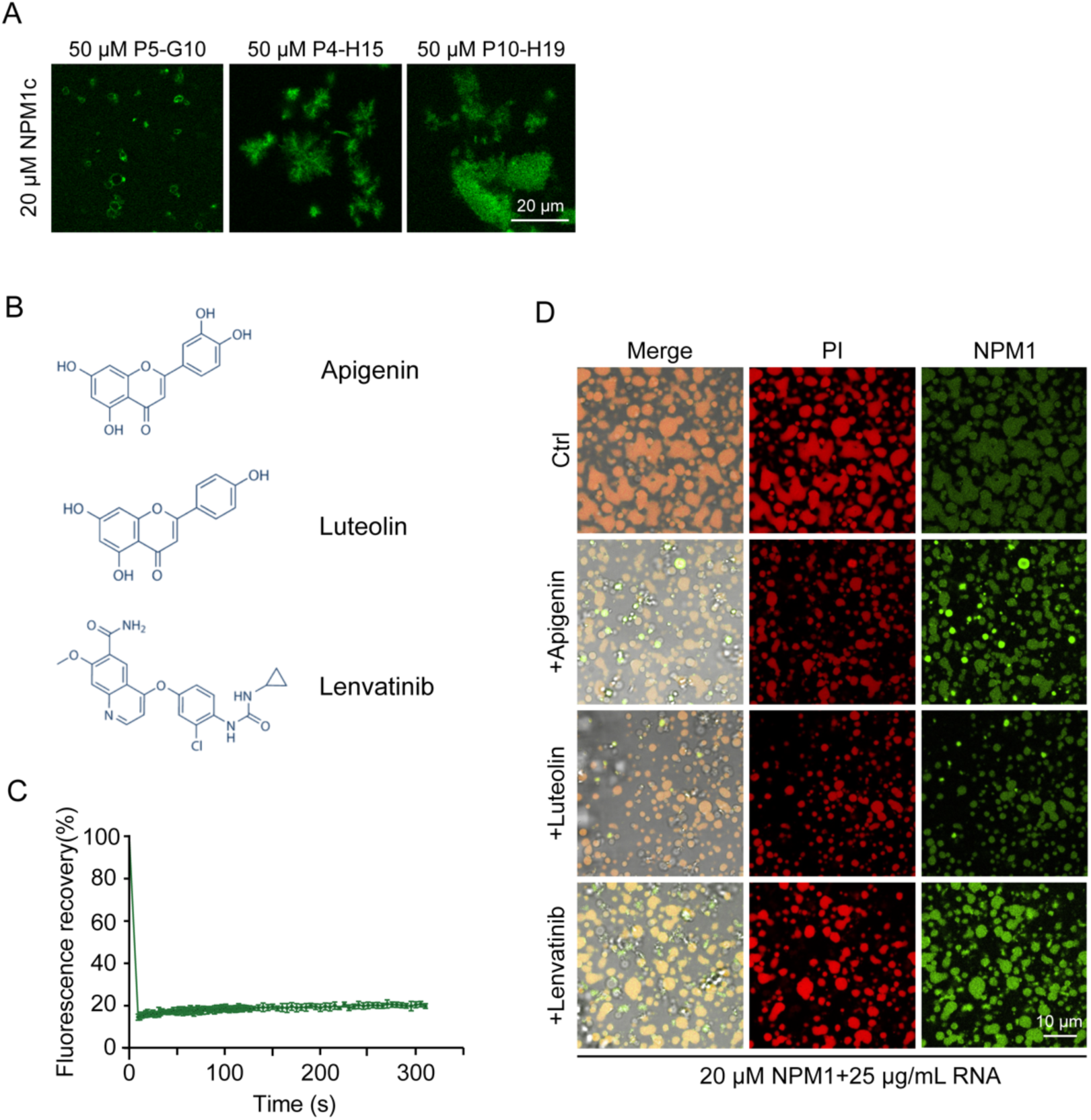
Characterization of compounds rescuing the impaired phase separation of NPM1c with RNA. **(A)** Representative images of NPM1c condensates induced by compounds in the presence of RNA (25 µg/mL) *in vitro*. **(B)** Molecular structures of Apigenin, Luteolin and Lenvatinib. **(C)** FRAP analysis of the dynamic behavior of NPM1c condensates in the presence of Lenvatinib and RNA *in vitro*. NPM1c, X 20M; RNA 25 µg/mL; Lenvatinib, 50 µM. **(D)** Representative images show the effects of compounds (50 µM) on NPM1/RNA condensates *in vitro*. NPM1 is labeled with Alexa488; RNA is labeled with PI.

**FigureS5.**
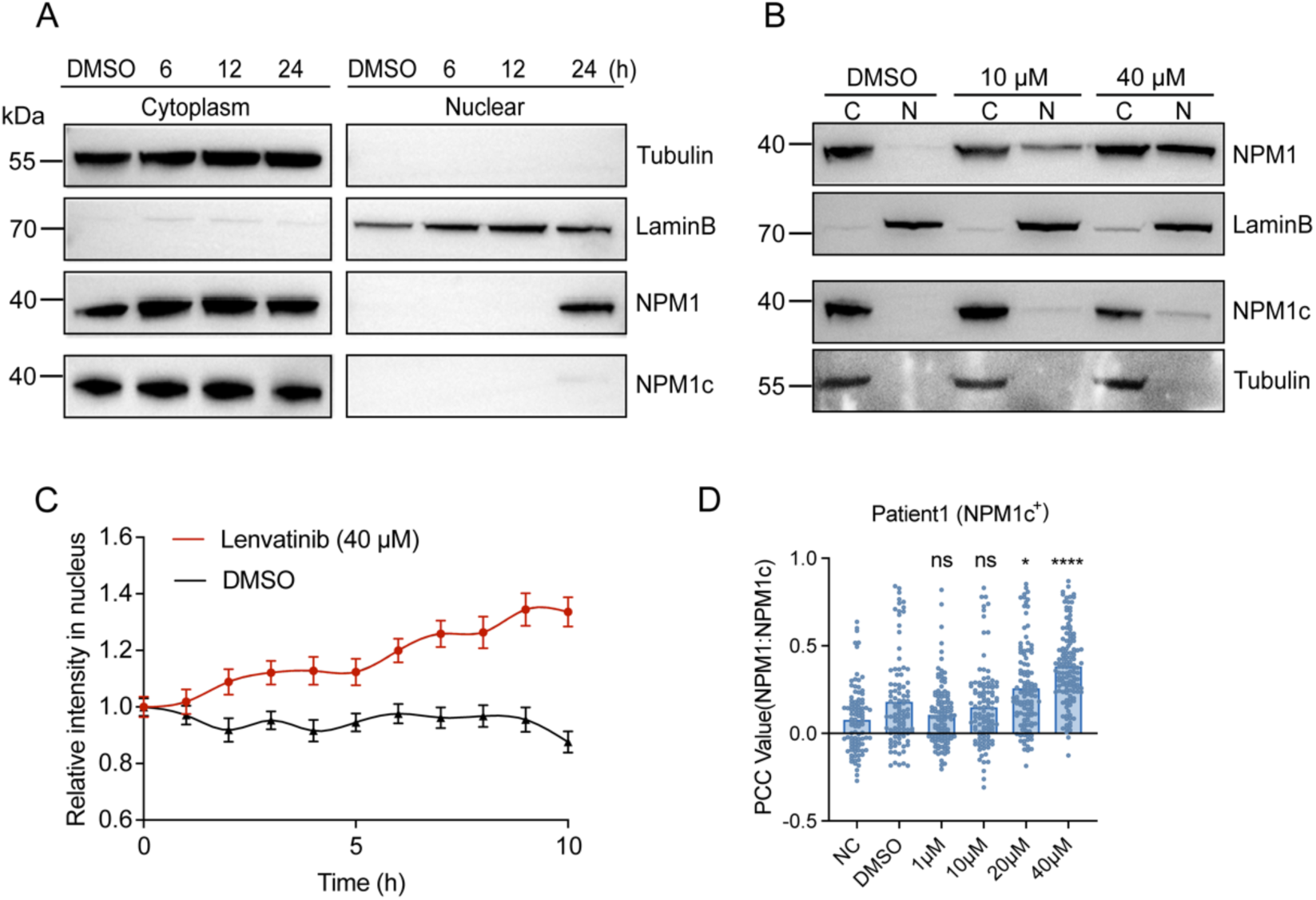
Lenvatinib reverses NPM1c mislocalization and exhibits anti-leukemia activity. (A-B) Western blot analysis of NPM1 and NPM1c protein levels in the cytoplasm and the nucleus of OCI-AML3 cells after exposure to Lenvatinib for different times (A) and at different concentrations (B). **(C)** Statistical analysis of the relative fluorescence intensity of mEGFP-NPM1c in the nucleus of HEK293T cells upon Lenvatinib treatment. **(D)** Statistical analysis of the co-localization value of NPM1/NPM1c in patient-derived blasts harboring a heterozygous *NPM1c* mutation. Cells were treated with Lenvatinib at the indicated concentrations for 24 hours.

## Experimental Procedures

### Cell culture

Human cancer cell lines, including HEK293T and Hela cells were cultured in DMEM (HyClone, Cytiva) plus 10% fetal bovine serum (GIBCO, Thermo Fisher Scientific), and 100 Units/ml Pen-Strep (GIBCO, Thermo Fisher Scientific), at 37°C, 5% CO2. HL-60 and Jurkat cells were cultured in RPMI-1640 (GIBCO, Thermo Fisher Scientific) plus 10% fetal bovine serum, and 100 Units /ml Pen-Strep, at 37°C, 5% CO_2_. OCI-AML3 cells were cultured in RPMI-1640 (GIBCO, Thermo Fisher Scientific) plus 20% fetal bovine serum, and 100 Units /ml Pen-Strep, at 37°C, 5% CO_2_.

NPM1c positive AML cells were obtained from patients at Peking University People’s Hospital. The patient has signed a written informed consent. Patient derived PBMC were extracted using Lymphoprep^TM^ (STEMCELL) and maintained in RPMI-1640 plus 20% fetal bovine serum, and 100 Units/ml Pen-Strep, at 37°C, 5% CO_2_.

### Protein expression and purification

The NPM1, NPM1c and NPM1c^288/290W^ constructs were cloned into the pRSFDuet-1 vector (Novagen, Merck), with an N-terminal poly-His tag. The recombinant proteins were overexpressed in *E. coli* BL21 (DE3). After overnight induction by 0.5 mM isopropyl β-D-thioacetamide (IPTG) at 16°C in LB medium, cells were harvested and suspended in the buffer: 40mM Tris-HCl (pH 7.4), 500mM NaCl, and 10mM imidazole. After cell lysis and centrifugation, the recombined proteins were purified with the HisTrap column and eluted with a linear imidazole gradient from 20 mM to 500 mM. The proteins were further purified through ion exchange column, and size-exclusion chromatography using a Superdex 200 Increase 10/300 GL column (Cytiva) in elution buffer (20 mM HEPES, pH7.4, 500mM NaCl).

### Protein labeling

Recombinant proteins were labeled with Alexa Fluor™ 647 C_2_ Maleimide or 488 C_5_ Maleimide (Thermo Fisher Scientific) according to the manufacturer’s protocol. In brief, proteins were incubated with dyes for 1 h at room temperature with continuous stirring. The free dyes were removed by centrifugation in Zeba^TM^ Spin Desalting Columns (Thermo Fisher Scientific, 89882).

### *In vitro* liquid-liquid phase separation (LLPS) assay

*In vitro* LLPS experiments were conducted at room temperature. For LLPS assays with RNA, recombinant proteins and RNA were diluted to the indicated final concentrations in the reaction buffer (20 mM HEPS, pH 7.4, 150 mM NaCl) with a total volume of 10 µl to induce phase separation. For LLPS assays with RnaseA treatment, a final concentration of 100 μg/mL RnaseA was add into pre-formed NPM1/RNA phase separation system containing 10 μM NPM1. The process of droplet reduction was recorded by continuous imaging.

For LLPS assays with PEG8000, recombinant proteins were diluted to the indicated final concentrations in the reaction buffer (20 mM HEPS, pH 7.4, 150 mM NaCl, 5% PEG8000) with a total volume of 10 µl to induce phase separation.

For LLPS assays with compounds, recombinant proteins and compounds were diluted to the indicated final concentrations in the reaction buffer (20 mM HEPS, pH 7.4, 150 mM NaCl) with a total volume of 10 µl to induce phase separation. Imaging was performed with a NIKON A1R HD25 confocal microscopy.

### Fluorescence Recovery After Photobleaching (FRAP) assays

The FRAP assay was conducted using the FRAP module of the NIKON A1R HD25 confocal microscopy system. Bleaching was focused on a circular region of interest (ROI) using 100% laser power, and time-lapse images were collected. Images were captured at the indicated time points to observe the dynamic changes of fluorescence intensity. Background intensity was subtracted and values are reported relative to pre-bleaching time points.

### Immunofluorescence assays

Cells were fixed in 4% paraformaldehyde for 15 minutes and permeabilized with 0.1% TritonX-100 in TBST. Subsequently, cells were blocked in 0.5% BSA (in PBS), and incubated with primary antibodies overnight at 4°C, washed 3 times with TBST, and incubated with secondary antibodies for 1 hour. The cells were then sealed in 4′,6′-diamidino-2-phenylindole (DAPI) (Beyotime), and examined with a NIKON A1R HD25 confocal microscope.

### RnaseA digestion in HEK293T cells

Briefly, HEK293T cells seeded on coverslips were treated with 0.5% Triton X-100 in CSL buffer (100 mM NaCl, 10 mM Pipes, 300 mM Sucrose, 3 mM MgCl2, 1 mM EGTA and cocktail) for 10 minutes at 4°C. Subsequently, cells were incubated with RNase A (10 μg/ml) in digesting buffer (100 mM NaCl, 10 mM Pipes, 300 mM Sucrose, 3 mM MgCl2, 1 mM EGTA and cocktail) in 37°C for 10 minutes. After washing the cells with PBS for three times, immunofluorescence was performed to confirm protein localization. Primary antibodies used for immunoblotting: Mouse anti-NPM1 monoclonal antibody (Thermo Fisher Scientific).

### Immunoblot (IB) assays

Cell lysates were obtained using Minute^TM^ Total Protein Extraction Kit for Animal Cultured Cells/Tissues (Invent Biotech) or Nuclear-Cytoplasmic Extraction Kit (Invent Biotech) according to the manufacturer’s instructions. Protein concentration was determined using the BCA Protein Quantification Kit (Promega). Protein samples were fractionated by SDS-PAGE and transferred to a PVDF membrane. The membranes were incubated overnight with primary antibodies, washed 3 times with TBST, and incubated with the corresponding secondary antibody at room for 2 hours. Finally, enhanced chemiluminescence reagent (Thermo Fisher) was used to visualize the results. Primary antibodies used for immunoblotting: Rabbit anti-NPM1c polyclonal antibody (Thermo Fisher Scientific), Mouse anti-NPM1 monoclonal antibody (Thermo Fisher Scientific), Mouse anti-β-Tubulin monoclonal antibody (Abmart), Rabbit anti-Lamin B polyclonal antibody (Proteintech).

### Immunoelectron microscopy and in situ hybridization

Lowicryl HM20-embedded thin sections were first blocked with 2% serum albumin in 100 mM (pH 7.4) phosphate buffer for 15 minutes, followed by incubation with primary antibody diluted in blocking buffer for 1.5 hours (dilution ratio is 1:5 for anti-NPM1c antibody). Subsequently, grids were washed with phosphate buffer for five times (2 min each) and incubated with the goat anti-mouse antibody (coupled with gold particle, dilution ratio is 1:50) for 1 hour. After washing with Tris–HCl (pH 7.4) buffer containing 1% serum albumin, grids were fixed with 1% glutaraldehyde. Subsequently, grids were washed with ddH_2_O for five times and contrasted for 5 min in 5% uranyl acetate. The stained sample was imaged in a Hitachi H-7650 transmission electron microscope (Hitachi) using an accelerating voltage of 80 kV.

### Immunoprecipitation assay

HEK293T cells were co-transfected with mCherry-NPM1 and mEGFP-NPM1c using Lipo8000 (Beyotime), following the manufacturer’s instructions. After 48 hours, cells were washed with cold PBS and then lysed by 500 μL ice-cold lysis buffer (Invent) containing protease inhibitor cocktail. One tenth of the lysate was taken out for a reference of input. The remaining lysate was incubated with GFP beads on a rotating platform for 1 hour at 4 °C. Next, the beads were washed for four times with wash buffer (PBS plus protease inhibitor cocktail) at 4 °C. The supernatant was then discarded, and the beads were resuspended with 70 μL 1× SDS loading buffer. The samples were then subjected to western blot analysis.

### Estimation of endogenous NPM1/NPM1c protein concentrations

Briefly, the quantification was based on the western blot analysis performed on cell lysates and purified NPM1/ NPM1c protein. Cell lysates of OCI-AML3 and HL-60 cells were subjected to western blot with 25-240 ng purified NPM1/ NPM1c protein. Standard curves were plotted using the gray values of purified recombinant NPM1 and NPM1c proteins with ImageJ. The volume of collected cells was calculated using the radius measured by confocal microscopy.

### Differential Scanning Calorimetry (DSC) Assay

DSC experiments were performed using a MicroCal PEAQ-DSC, with approximately 1mg/mL of each testing sample s used for the experiments. The equipment was first washed with 50 mL ddH_2_O. 250 μL proteins were then added after 2 cycles of buffer-buffer scan. Samples were heated at 1 °C/min from 50 °C to a maximum temperature of 130 °C. Thermal curves, as well as the Tm and Cp values, were calculated by the software.

### *In vitro* centrifugation-based phase separation assay

The purified NPM1, NPM1c and NPM1c^288/290W^ were centrifugated at 15000 rpm for 10 minutes to remove aggregates. For single component protein detection, final concentration of 20 μM proteins and 100 μg/mL RNA were mixed at a volume of 100 μL and incubated at 4 °C overnight. For two component protein detection, NPM1 and NPM1c were mixed at a final concentration of 10 μM and incubated at 4 °C overnight. Then, samples were centrifuged at 15000 rpm for 30 min at room temperature. Supernatant and pellet were separated into two tubes immediately after centrifugation. The pellet fraction was washed with the LLPS buffer and re-suspended with the same buffer to the equal volume as supernatant. Proteins from supernatant and pellet fraction were separated by 4-20% SDS-PAGE and the gel was stained with Coomassie blue.

### Flow cytometry-based differentiation marker detection

Cells (1×106) were treated with Lenvatinib at the indicated concentrations for 96 hours. Subsequently, cells were stained with PE-CD11b (Biolegend) and PE-CD14 (Biolegend) antibodies according to manufacturer’s instruction. Flow cytometry was performed to measure the expression levels of CD11b and CD14.

### Cell Viability Assay

Cells were seeded in 96-well plates and exposed to Lenvatinib at the indicated concentrations for 24, 48, 72 and 96 hours. The CCK8 cell viability assay was performed to assess the cell viability.

### Flow cytometry-based apoptosis detection

Cells (1×10^6^) were treated with Lenvatinib at the indicated concentrations for 24, 48, 72 and 96 hours. Cells were then harvested and stained with Annexin V-FITC and Propidium Iodide (Yeason) for 15 minutes at room temperature. Measurements were taken on BD LSRFortessa SORP.

### Flow cytometry-based cell cycle detection

Cells (1×106) were treated with Lenvatinib at the indicated concentration for 24 hours. Subsequently, cells were washed twice with cold PBS and fixed in 75% ethyl alcohol at 4 °C overnight. After washing with cold PBS, cells were stained in PBS containing 50 μg/mL PI and 100 μg/mL RnaseA. Measurements were taken on BD LSRFortessa SORP.

### Data analysis

Images were processed with NIS-Elements AR Analysis (Nikon, Inc.) and Adobe Illustrator CC (Adobe Systems, Inc.). The fluorescence intensity was analyzed using Image J (National Institutes of Health) and the corresponding graphs were generated by GraphPad Prism 8.

